# GWANN: Implementing deep learning in genome wide association studies

**DOI:** 10.1101/2022.06.01.494275

**Authors:** Nimrod Ashkenazy, Martin Feder, Ofer M. Shir, Sariel Hübner

**Affiliations:** Galilee Research Institute (MIGAL), Tel-Hai Academic College, Upper Galilee, 11016, Israel

## Abstract

**Motivation:** Genome wide association studies (GWAS) are extensively used across species to identify genes that underlie important traits. Most GWAS methods apply modifications and extensions to a linear regression model in order to detect significant associations between genetic variation and a trait. Despite their popularity, these statistical models tend to suffer from high false positive rates, especially when utilized on large variant datasets or complex demographic scenarios. To overcome this, aggressive statistical corrections are applied which frequently diminish true associations.

**Results:** Here we consider a deep learning approach, and present an implementation of a convolutional neural network (CNN) to identify genetic variation that is associated with a trait of interest. To exploit the strength of CNNs in visual recognition, the genotype information is represented as an image, which enables the model to correctly classify genetic variants with respect to the trait, even when a population structure is present. Our proposed approach was implemented in a package called GWANN which exhibited solid performance. Overall, GWANN outperformed popular GWAS tools on both simulated and real datasets, and enabled the identification of association signals with increased sensitivity and speed.

**Availability and implementation:** The package is available at: https://github.com/hubner-lab/GWANN

## Introduction

Identification of genomic regions and candidate genes that are associated with a trait of interest is pivotal for understanding the genetic mechanism that underlies an observed diversity. To address this, a plethora of tools were developed over the years, implementing different approaches adjusted to fit the target trait and the studied population (Uffelmann et al. 2021). Among these approaches, genome wide association studies (GWAS) have gained enormous popularity and are extensively used to study different species, traits and data types, owing to their generality, flexibility and simplicity of implementation (Visscher et al. 2017, Cortes et al. 2020). Most GWAS tools employ a linear model to estimate the contribution of a genetic marker (e.g. single nucleotide polymorphism – SNP) to a trait which is used as the response variable in the model. This approach was further extended to allow the inclusion of additional factors in the model which are essential for controlling the rate of spurious signals. Despite the adjustments and improvements that were applied to the basic statistical model, this approach suffers from a high false positives rate, reflecting the difficulty to broadly control it due to specific constraints of population stratification, genetic relatedness, ascertainment bias and so forth (Korte and Farlow 2013). Moreover, the statistical test is independently conducted for each SNP along hundreds of thousands or millions of variants, leading to a high number of independent tests, thus requiring intensive correction for multiple comparisons. These corrections are often too stringent and lead to an increase in the false negative rate to the detriment of the study objectives (Brzisky et al. 2017, Chen et al. 2021). Therefore, an analytical approach that is less prone to statistical constraints can potentially obtain a clearer signal for associations between genetic markers and a trait of interest.

Recent developments in Machine Learning (ML) algorithms, and specifically artificial neural networks (ANN), have led to an overwhelming success in tackling computational tasks in image and sound recognition, and other complex classification problems (LeCun et al. 2015, Webb 2018). Among ANN models, convolutional neural networks (CNN) have been highly successful in solving complex image recognition problems, at prominent success rates that often exceed human performance (Li et al. 2021). In short, ML algorithms typically compute the feature-space and its mapping to the target value in a supervised-learning fashion. That is, supervised-learning aims to obtain a model that returns a prediction, or a classifier, for a so-called target value when given labeled features as input. Modern ANN models have grown to become large networks, entitled Deep Neural Networks, leading to the coined terminology Deep Learning (DL).

Overall, the latest ML and DL developments have also contributed to practical problem-solving in the broad domain of genomics (Mazurenko et al. 2020). Explicitly, CNNs are increasingly being implemented to address complex problems in genomics including identification of RNA and DNA binding sites (Alipanahi et al. 2015), modeling regulatory elements (Zhou et al. 2015), genetic variant calling (Poplin et al. 2018), and prediction of phenotypes (Liu et al. 2019). Despite their efficiency, implementations of DL models in genomic mapping problems are largely lacking.

Here, we present an implementation of a DL approach using CNN to identify genomic polymorphisms that are associated with a quantitative trait of interest. To address this, simulated genomic data was represented as a sorted image in accordance with a simulated trait distribution, and was used to train the CNN model. To correct for potential high false positive rates imposed by strong population structure, a set of true negative images were also incorporated during the training phase of the model. Following this, real-world data was analyzed to identify signals of associations between genetic variants and an observed trait in a diverse sunflower population.

## Materials and Methods

### Data structure and conversion of genomic data to image

Standard genomic variants data is stored in a matrix-like format such as variant call format (VCF) file where individuals are presented in columns and genomic polymorphism information is indicated in rows. For each polymorphic site (e.g. SNP) genotyped in each diploid individual, 1 out of 4 features could be assigned in agreement with 3 possible genotypes and missing data: homozygote for the minor alleles (0), homozygote for the major allele (2), heterozygote (1) and missing data (3). Each polymorphic site is characterized across all accessions so *v*_*i*_ ∈ [0,1,2,3] and the length of *v* equals the number of individuals. Each SNP is independently tested for association with the measured trait, thus an image is produced for each SNP across all individuals from the information stored in *v*.

Encoding the SNP data into an image is conducted after sorting the vector *v* in accordance with the trait scores, namely individuals with low scores are at the top. The vector is then folded into a matrix with predefined number of rows (e.g. 6 individuals into a matrix with 3 columns):

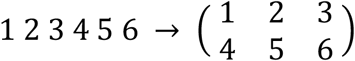

The constructed images (matrices), each representing a SNP, are classified as either associated or not-associated with the trait (i.e., binary labeling) by a CNN implemented in `python3` using the PYTORCH package (Paszke et al. 2019). The CNN model was chosen as the learning algorithm because it has the flexibility to identify patterns despite substantial noise, which is expected in genetic data obtained from natural populations. Explicitly, the network is comprised of 3 convolutional layers and 2 max-pooling layers to introduce non-linearity, and 2 fully-connected linear layers. The output of the network was classified as positive (0 inclusive) if a SNP was identified as associated, and negative for not-associated SNP.

To quantify the performance of the network, the standard ML success-rates were utilized, using the following forms:

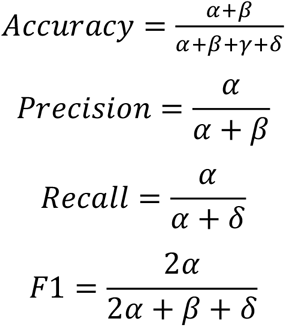

Where *α* denotes the number of true positives, *β* corresponds to false positives, *γ* corresponds to true negatives, and *δ* corresponds to false negatives.

### Simulation and training the model

To train the model, genomic data is simulated for a population comprised of a user defined number of individuals in accordance with the size of the studied test population. Genomic data is simulated with the GENOME package (Liang et al. 2007), which implements a standard coalescence model to draw the population data. Trait scores were generated for the simulated population with PHENOSIM (Günther et al. 2011) based on GENOME simulations. The number of populations simulated to train the model is defined by the user and so are the number of SNPs genotyped in each population, the expected number of associated SNPs, the minor allele frequency and the missing data rate. We recommend to set these parameters in accordance with the expectations in the studied population. For testing the performance of GWANN, we simulated 100 populations, each comprised of 300 individuals that were genotyped at 10,000 sites (SNPs) of which 100 are associated with the trait with equal effect on the trait. The minor allele frequency was set to 5% and the rate of missing data to 3%. Following the genomic data simulation, a trait score was simulated for each individual and an image was generated for each SNP after sorting individuals by the trait score. The produced images were then used to train the model using a stochastic gradient descent as the solver (i.e., optimizer) and utilizing mean square error as the loss function. In some populations, a strong pedigree relationship among individuals may confound the model and increase the rate of false positive calls. To correct for potential population structure, simulations can be conducted in accordance with the number of sub-populations expected in the studied population. Correction for population structure is applied by generating sorted images in accordance with the genetic similarity among individuals and using these images as true negatives in the training phase. Genetic similarity is calculated as the level of identity by state between individuals and converted to a vector using multidimensional scaling (MDS). After training the model for a reasonable number of epochs (e.g. 500), the model assigns each SNP a two-valued output: the binary prediction, where values higher than 0 indicate a predicted association with the trait; and a prediction measure which is indicative for the certainty of the model.

### Testing GWANN with real data

To test the ability of GWANN to detect SNPs that are associated with a trait of interest in a real dataset we used the available data for the sunflower association mapping (SAM) population. The SAM population is comprised of 287 individuals that well represent the genetic variation among sunflower varieties and genetic and phenotypic data is available. Here, we used a publicly available data of approximately 600,000 SNPs with a maximum missing data rate of 30% (Hübner et al. 2019) and the branching phenotype scores (Mandel et al. 2013). This target trait was scored quantitatively and is highly important in sunflower breeding and allows to distinguish between breeding types. In addition, the SAM population is characterized by four major groups (Hübner et al. 2019), thus this population provides a good case-study for the performance of GWANN with real data. To compare the performance of GWANN to existing GWAS tools, the popular EMMAX package (Kang et al. 2010) was used to detect SNPs associated with branching in the same dataset.

## Results

To evaluate the performance of the GWANN application, various scenarios were simulated and tested. All runs were conducted on a personal computer with NVIDIA GeForce GTX 1080 Ti, a CUDA version 11.0, and CUDNN version 8005. Analyses were conducted in batches of 20 populations and 60 images (SNPs) per population at each round and performed very fast in practice. For example, training 1000 epochs for images of 20 columns and 15 rows (300 individuals) took approximately 30 seconds on this PC and was extended by additional 10 seconds when PYTORCH and CUDA were set to work deterministically (‘deterministic’ flag) to allow reproducibility across tests.

To identify SNPs that are associated with a trait of interest, GWANN follows three steps: 1) simulation of data and conversion of each SNP to an image, 2) training the model based on true positive and true negative data instances, and 3) prediction of association in a tested dataset. To evaluate the performance of GWANN under different image structures, namely the number of columns in the matrix and the minor allele frequency, we simulated one hundred populations comprised of 300 individuals each. First, simulated data was fixed with a minor allele frequency of 5% and 3% missing data, and the converted images were organized in an exceeding number of columns from 10 to 30. For each population, a total number of 10,000 SNPs was simulated, featuring a fixed number of 100 SNPs that are expected to be associated with the trait. The model was trained with 200 SNPs at each round, where 100 SNPs were assigned as true positives (associated) and 100 SNPs were assigned as true negatives (not-associated) in each population. Among the 100 populations simulated, 70 were used to train the model and 30 were used as test populations to classify SNPs.

Overall, the number of columns in the generated images (matrices) had a mild effect on the accuracy of the model, varying between 74-76% across different structures. The highest precision (93%) and F1 (78%) were obtained at 25 columns, thus an image structure of 25×12 was fixed to test the effect of minor allele frequency on the model performance (Figure S1).

To test the sensitivity of the model to different values of minor allele frequency (MAF), the analysis was performed for MAF values ranging from 5%-35% (Figure S2). The observed pattern of an associated SNP image dramatically varied in accordance with the minor allele frequency which may challenge the learnability of the model (Figure 1). As expected, the model was very sensitive to the value of minor allele frequency specifically at values below 5% where the model accuracy (74%), precision (82%), recall (75%) and F1 (75%) were lowest. However, when the minor allele frequency exceeded 5%, the accuracy (85%), and all other measures (precision = 95%, recall = 81%, F1 = 86%) quickly increased with little improvement at values higher than 10%. The peak performance was observed at MAF = 35% (accuracy = 95%, precision = 97%, recall = 90%, F1_MAF35%_ = 95%), where the pattern in the image was evidently very pronounced.

**Figure 1:**
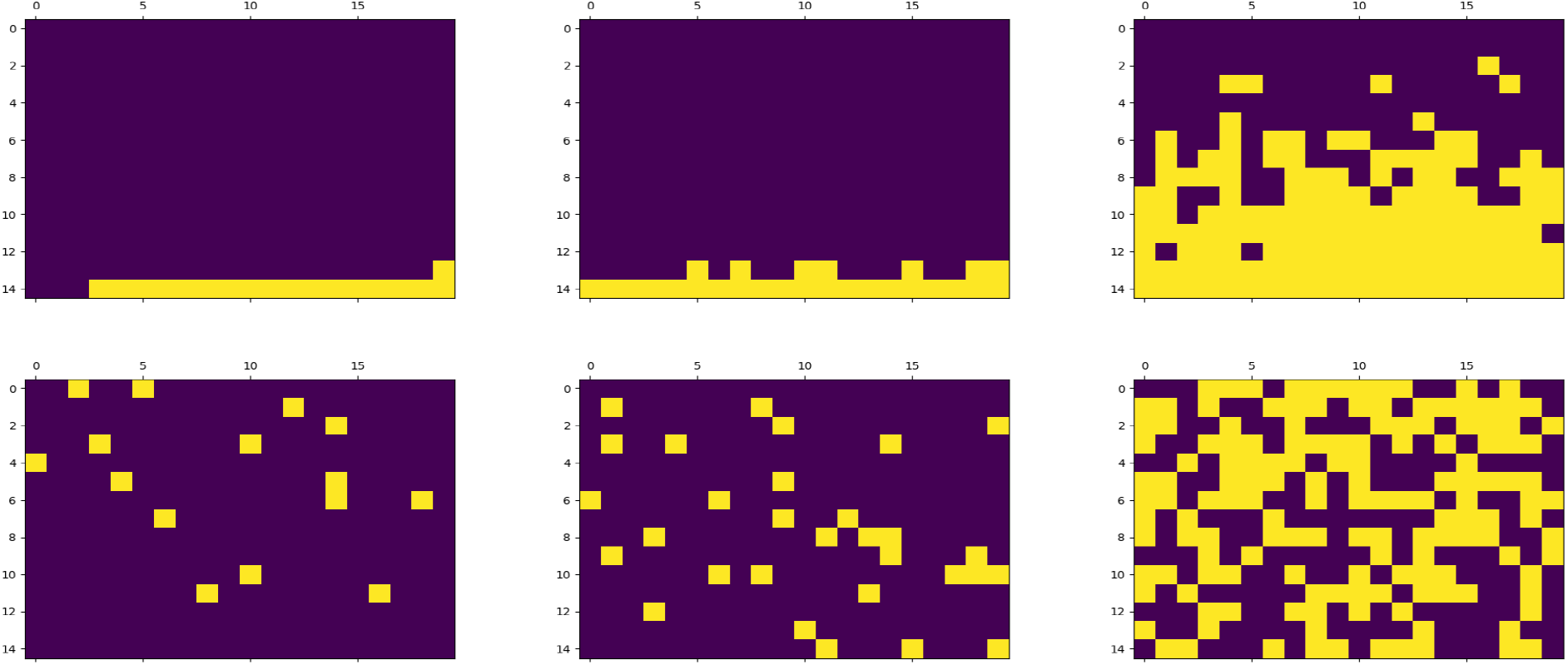
Encoding a single variant data into an image. At the top are images for three representing true associated SNPs and at the bottom are three random unassociated SNPs. Images are organized from left to right with increasing minor allele frequency of 5%, 10%, and 50%.

Next, the performance of GWANN was evaluated with real data obtained for the sunflower association mapping (SAM) population. This population is comprised of 287 individuals which could be divided into 4 groups in accordance with breeding background, thus a clear population structure is present among accessions. Genotypic data is publicly available for this population and include 600 thousand SNPs with minor allele frequency of 5% and maximum missing data rate of 30%. To train the model, 100 populations of 300 individuals each were simulated with minor allele frequency of 5%. For each population, 10,000 variants were simulated with a fixed rate of 1 associated call for every 50 non-associated variants. To compare the performance of GWANN to a standard mixed linear model application, the EMMAX program was used with default parameters and the same genotypic and phenotypic data. Population structure was corrected in EMMAX as covariates using the 4 first PCs from a PCA conducted for this population and the significance threshold was set at 10^−7^ after correcting for multiple-testing using the simpleM algorithm (Gao et al. 2008). Both EMMAX and GWANN were able to detect a strong signal on chromosome 10 confirming previous studies conducted for this population and trait (Mandel et al 2013). However, GWANN was able to detect additional signals on chromosomes 13 and 8 (Figure 2, Figure S3) which were previously identified in the context of branching (difference between female and restorer lines in sunflower) using genome scans approach (Hübner et al. 2019).

**Figure 2:**
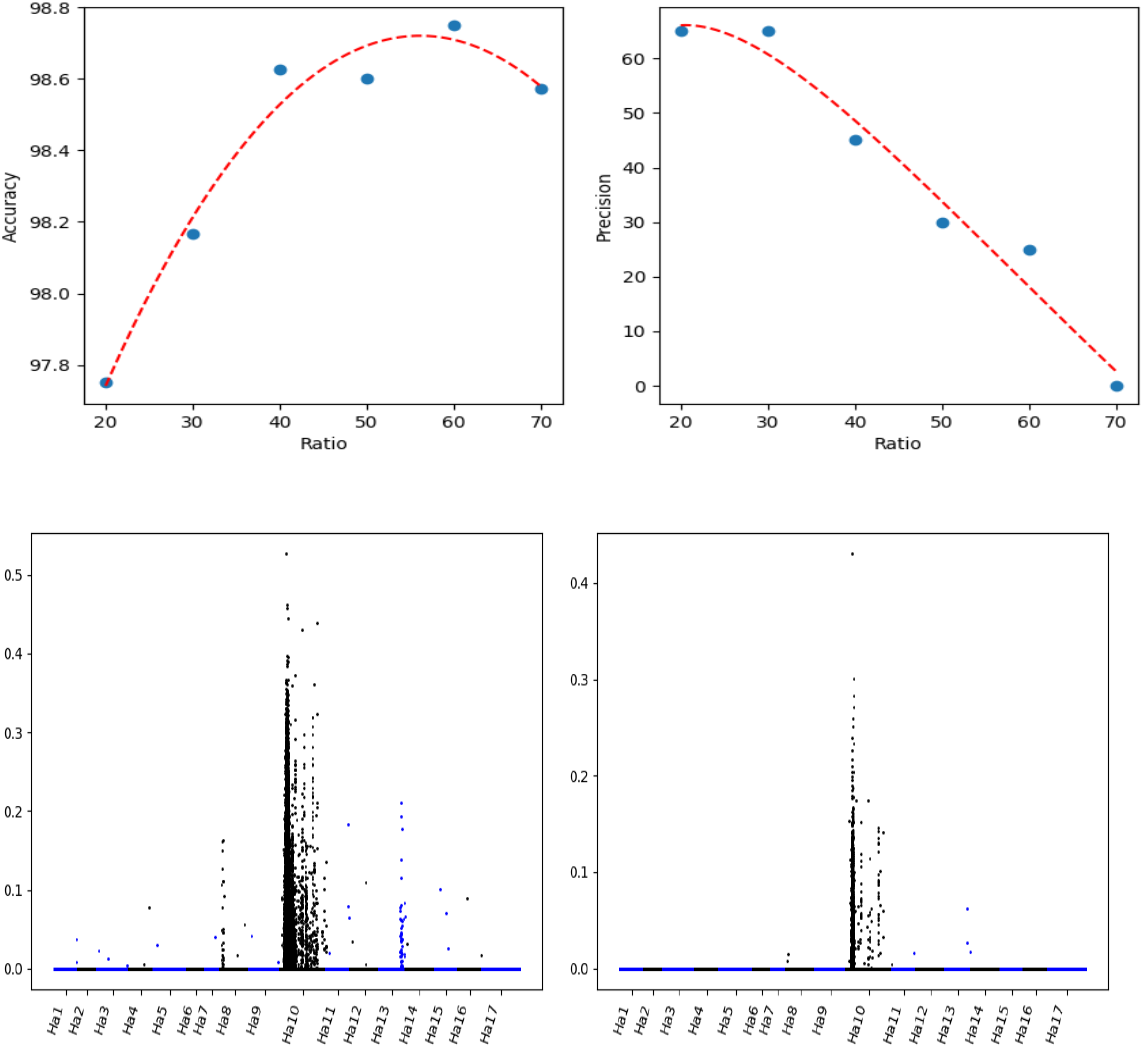
Evaluation of GWANN performance with the branching trait in sunflower. Testing the effect of associated SNP sampling rate in the training dataset on the detection power of GWANN. In the upper panel are accuracy (left) and precision (right) for increasing rate of expected associated SNPs. In the lower panel are Mnahattan plots generated for a sampling rate of 1:50 (left) and 1:70 (right) associated:not-associated SNPs

Sensitivity of the model is largely affected by the sampling rate of associated SNPs in the training populations, namely higher rate of SNPs that are expected as associated with the trait will increase the sensitivity of the model. To address this, different sampling rates between 0.001 and 0.05 were tested and confirmed that higher sampling increases the number of SNPs classified as associated (Figure 2, Figure S4). This parameter should be adjusted by the user in accordance with the studied population and trait, thus it is recommended to explore GWANN with different sampling rates.

## Discussion

With the dramatic increase in sequencing capacity it is now possible to generate genomic data for large populations at a reasonable cost even for species with large or complex genome. This technological advent allows to study the genetic basis of important traits at the finest resolution, yet current statistical methods often suffer from increased false negative rate due to implementation of a stringent control for the false positive rate (Korte and Farlow 2013, Brzyski et al. 2017, Chen et al. 2021). This caveat in linear regression statistical models and related extensions is exacerbated with the increase in genotyping capacity and resolution. Here, we present a new Deep Learning (DL) approach to identify genetic variation that is associated with a measured trait of interest. A CNN model was chosen as the learning algorithm because it was shown to perform well in image recognition and classification even when the data was noisy (Li et al. 2021). To facilitate an efficient learning by the model, we developed a conversion method of genotype data to an image based on sorted information, thus the problem was reduced to a classification problem of an image pattern. Based on simulations and real data analysis, the most important parameter affecting the sensitivity of the model is the rate of expected associated SNPs in the entire dataset indicating that more sampling may increase the rate of false positive detection. Thus, it is strongly recommended to explore the rate of sampling for each population and phenotype and adjust it accordingly. Other parameters, including the image (matrix) structure and minor allele frequency had mild effects on the results except in extreme scenarios of unidimensional image or minor allele frequency below 5%.

The implementation of GWANN include a training phase of simulated data which reduce the overfit of the model when predicting in the test population. Despite this advantage, performance may deteriorate if the simulated population poorly represent the tested population, thus users should adjust the parameters, namely population size, minor allele frequency, population structure and expected sampling rate, to the studied population to guarantee highest performance. Overall, the algorithm implementation had higher sensitivity and speed compared with the popular mix linear model implemented in the EMMAX package. In GWANN, the entire process, including training and prediction, has a duration of less than 30 seconds on a standard PC, allowing the user to explore different parameters. The concept, as well as the implemented approach to convert genomic data into a learnable image, can be further improved to include more complex population scenarios, traits combinations and so forth.

